# Computational Chromosome Conformation Capture by Correlation of ChIP-seq at CTCF motifs

**DOI:** 10.1101/257584

**Authors:** Jonas Ibn-Salem, Miguel A. Andrade-Navarro

**Author notes:** corresponding authors: Jonas Ibn-Salem, Miguel A. Andrade-Navarro.

## Abstract

We present a computational method to gain knowledge of the three-dimensional structure of the genome from ChIP-seq datasets. While not designed to detect contacts, the ChIP-seq protocol cross-links proteins with each other and with DNA. Consequently, genomic regions that interact with the protein binding-site via chromatin looping are coimmunoprecipitated and sequenced. This produces minor ChIP-seq signals around CTCF motif pairs at loop anchor regions. Together with genomic sequence features, these signals predict whether loop anchors interact or not. Our method, Computational Chromosome Conformation Capture by Correlation of ChIP-seq at CTCF motifs (7C), is available as an R/Bioconductor package: http://bioconductor.org/packages/sevenC

## Background

The three-dimensional folding structure of the genome and its dynamic changes play a very important role in the regulation of gene expression [1–3]. For example, while it was well known that transcription factors (TFs) can regulate genes by binding to their adjacent promoters, many TF binding sites are in distal regulatory regions, such as enhancers, that are hundreds of kilo bases far from gene promoters [4]. These distal regulatory regions can physically interact with promoters of regulated genes by chromatin looping interactions [5–7], thus it is not trivial to associate TFs to regulated genes without information of the genome structure [8]. Such looping interactions can be measured by chromosome conformation capture (3C) experiments [9] and its variations to either study all interactions from single targeted regions (4C) [10] or multiple target regions (5C) [11], interactions between all regions genome-wide (Hi-C) [12,13] or interactions mediated by specific proteins (6C [14] ChIA-PET [15,16], and HiChIP [17]).

While these experimental methods have brought many exciting insights into the three-dimensional organization of genomes [1–3,18], these methods are not only elaborate and expensive but also require large amounts of sample material or have limited resolution [19,20]. As a consequence, genome-wide chromatin interaction maps are only available for a limited number of cell types and conditions.

In contrast, the binding sites of TFs can be detected genome-wide by ChIP-seq experiments, and are available for hundreds of TFs in many cell types and conditions [21–23]. Here, we propose that it is possible to use these data to detect chromatin loops.

Recent studies provide functional insights about how chromatin loops are formed and highlight the role of architectural proteins such as CTCF and cohesin [1]. CTCF recognizes a specific sequence motif, to which it binds with high affinity [24,25]. Interestingly, CTCF motifs are present in convergent orientation at chromatin loop anchors [13,16,26]. Furthermore, experimental inversion of the motif results in changes of loop formation and altered gene expression [27–29]. Polymer simulations and experimental perturbations led to a model of loop extrusion, in which loop-extruding factors, such as cohesin, form progressively larger loops but stall at CTCF binding sites in convergent orientation [29–31]. According to these models, CTCF binding sites can function as anchors of chromatin loops.

Our hypothesis is, that we can use convergently aligned CTCF motifs to search for similar ChIP-seq signals at both sites of chromatin loops to predict looping interactions from the largely available ChIP-seq data in many diverse cell-types and conditions (Fig. 1A). We then developed and tested a computational method to predict chromatin looping interactions from only genomic sequence features and TF binding data from single ChIP-seq experiments. We show that our method has high prediction performance when compared to Hi-C and ChIA-PET loops and that prediction performance depends on the ChIP-seq target, which allows screening for TFs with potential novel functions in chromatin loop formation. The predicted looping interactions can be used to (I) associate TF binding sites or enhancers to regulated genes for conditions where Hi-C like data is not available, and (ii) to increase the resolution of interaction maps, where low resolution Hi-C data is available. We implemented our method as the R package *sevenC*.

**Fig. 1.**
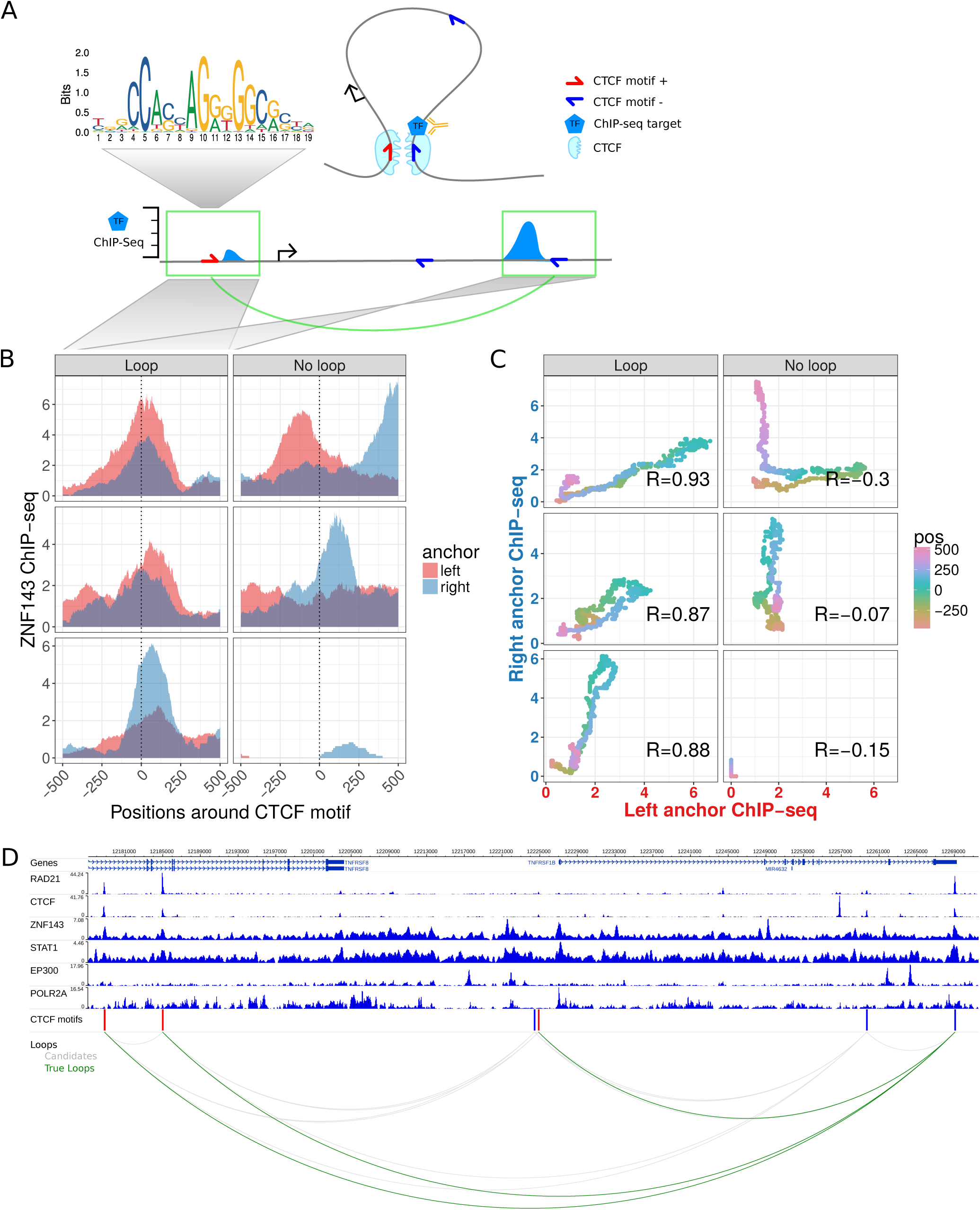
Chromatin looping interactions result in ChIP-seq coverage signals at direct and indirect bound loop anchors. **(A)** Schematic illustration of a chromatin loop with CTCF motifs at the loop anchors (top right). A transcription factor (TF) binds directly at the right loop anchor close to the CTCF motif. This results in a ChIP-seq coverage peak at the directly bound locus (bottom right) and in a minor signal at the other loop anchor (bottom left), both at the same distance to each CTCF motif. **(B)** Znf143 ChIP-seq coverage at six selected example CTCF motif pairs of which the ones in the left panel interact via loops according to Hi-C and ChIA-PET data and the ones in the right panel do not interact. The ChIP-seq coverage signal for each loci pair is shown in red for the left anchor region and in blue for the right anchor region, according to the distance to the CTCF motif (x-axis). Interacting CTCF motif pairs show more similar ChIP-seq coverage signals, which are often enriched at similar distances to the CTCF motif pairs, while the profiles of non-interacting pairs are less similar. **(C)** The similarity of ChIP-seq profiles by correlation of the ChIP-seq coverage signals of the selected motif pairs in (B). For each pair, the coverage at the right anchor is plotted versus the coverage at the left anchor at the same distance (color coded) from each CTCF motif. The Pearson correlation coefficient (R) of the dots is higher for interacting loci pairs. **(D)** Example loci on chromosome 1 shown in the genome-browser with six ChIP-seq tracks. Red and blue bars indicate CTCF recognition motifs on the forward and reverse strand, respectively. The bottom panel shows CTCF motif pairs in gray (candidates) and actually interacting pairs in green, according to ChIA-PET and Hi-C data.

## Results

### CTCF motif pairs as candidate chromatin loop anchors

In order to predict chromatin looping interactions from ChIP-seq data, we first analyzed which features at looping anchors correlate with interaction signals. As a starting point for all analyses we used 38,316 CTCF motif sites in the human genome as potential chromatin loop anchors. We built a dataset of all CTCF motif pairs located within a genomic distance of 1 Mb to each other. This resulted in 717,137 potential looping interactions; we expect that only a minority of these motif pairs will be in physical contact for a given cell type and condition. To label motif pairs as true loops, we used chromatin loops from published high-resolution *in-situ* Hi-C data and ChIA-PET data for CTCF and Pol2 in human GM12878 cells [13,16]. If a motif pair was measured to interact in one of the data sets, we labeled it as true interaction (Fig. S1). Overall 30,025 (4.19 %) of CTCF motif pairs were considered as true loops using these data sets.

### Similarity of ChIP-seq signals at looping CTCF motifs

The ChIP-seq protocol involves a cross-linking step, in which formaldehyde treatment results in covalent bonds between DNA and proteins [32]. This allows the pull-down and detection of sites directly bound by the targeted protein. However, cross-linking occurs also between proteins, which results in detection of sites that are indirectly bound through protein-protein interactions or chromatin looping interactions [33,34].

We hypothesized that if a protein binds directly to a genomic region in chromatin contact with other genomic regions, DNA from both loci might be pulled out in the cross-linking and DNA-purification step of ChIP-seq protocols. As a result, we expect ChIP-seq signals (e.g. mapped reads) at both genomic regions: the directly bound one and the chromatin loop interaction partner locus (Fig. 1A). Some proteins, like CTCF and potentially also RAD21, might act as homo-dimer at loop anchors. If both loop anchors are bound directly by dimerizing proteins, we expect the ChIP-seq signal at a similar distance to the CTCF motif. Thereby we assume the loop forming complex to be symmetric, that is, that the distance of the direct binding site to the CTCF motif center is the same on both anchors. For cohesin, for example, it was shown that it binds slightly upstream of the CTCF motif [16].

To test our hypothesis, we used CTCF motif pairs as anchors and compared the ChIP-seq signal from one anchor to the (reversed) signal of the corresponding anchor. We found similar ChIP-seq coverage patterns around CTCF motifs more often when the two sites perform looping interactions than when they do not (Fig. 1B). To quantify the similarity of ChIP-seq coverage from any two CTCF sites, we correlated their ChIP-seq signals at ±500 bp around the CTCF motif (Fig. 1C) (see Methods for details). Measuring ChIP-seq profile similarity by correlation has the advantage that the correlation can be high even if the anchor that is not bound directly has a much lower ChIP-seq signal (which is often the case).

Next, we compared ChIP-seq similarity at looping and non-looping CTCF motif pairs for six selected TF ChIP-seq data sets (Fig. 2A). Compared to non-interacting CTCF sites the ChIP-seq correlation is significantly higher at looping interactions (Fig. 2A). However, the overall correlation as well as the difference between looping and non-looping CTCF sites varies between TF ChIP-seq datasets (Fig. 2A). As expected, we observed a large difference for the CTCF ChIP-seq dataset but, interestingly, also for other known architectural proteins, such as Rad21 and Znf143. Moreover, other TFs, such as STAT1 have significantly higher ChIP-seq signal similarity at CTCF motifs that interact via chromatin looping. Overall, this analysis shows that ChIP-seq signals are more similar at interacting CTCF sites, indicating that this similarity can be used to predict looping interactions.

**Fig. 2.**
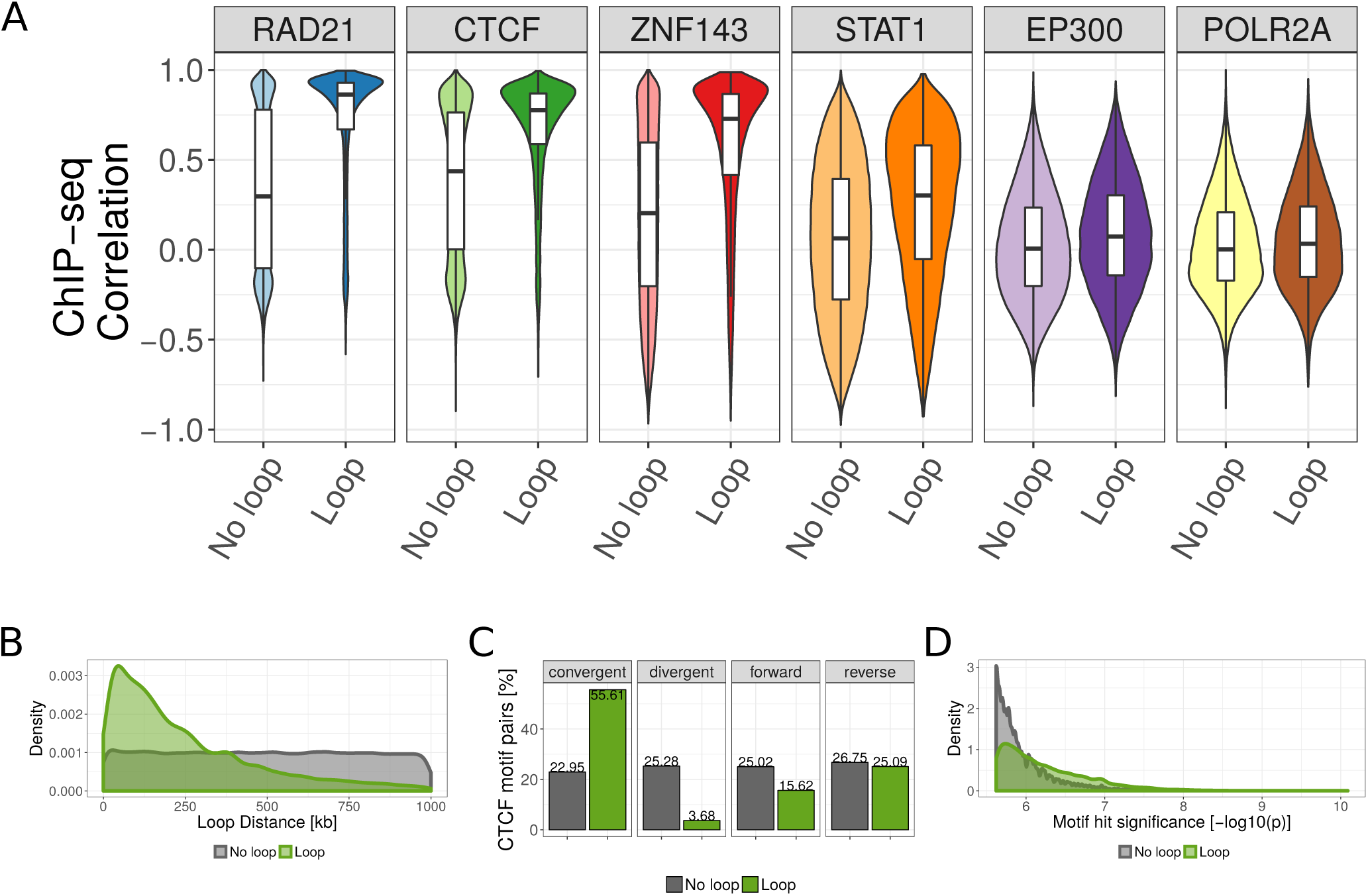
ChIP-seq similarity and genomic features of looping and non-looping CTCF motif pairs. **(A)** Boxplot of Pearson correlation coefficient of ChIP-seq signals between CTCF motif pairs for all CTCF motif pairs within 1 Mb genome-wide. The correlation is shown separately for non-looping and looping motif pairs (according to HI-C and ChIA-PET data in GM12878 cells), and for six selected ChIP-seq data sets in GM12878 cells. **(B)** Distance distribution between looping (green) and non-looping CTCF motif pairs. **(C)** Percent of looping and non-looping CTCF motif pairs in convergent, divergent, both forward, or both reverse orientation. **(D)** Distribution of CTCF motif hit significance as −log10 transformed p-value for looping and non-looping CTCF motif pairs. For each motif pair only the less significant motif is considered.

### Genomic sequence features of CTCF motif pairs are associated with looping

The frequency of two genomic regions to physically interact depends on their genomic distance [12]. Consequently, we observed that CTCF motif pairs are more often in contact when they are close to each other in the genomic sequence (Fig. 2B). Recent studies on 3D chromatin structure led to an increased understanding of the molecular mechanism of chromatin loop formation and suggested a functional role of CTCF proteins, which bind specific DNA sequences [1]. The canonical CTCF motif is non-palindromic and therefore occurs either in the positive or in the negative DNA strand. Importantly, it is known that CTCF motifs occur predominantly in convergent orientation to each other at chromatin loop anchors [13,26]. Experimental inversions of CTCF motifs lead to changes of the interactions and expression of the associated genes [27,28]. Accordingly, we observed that 55.6% of the looping CTCF pairs have convergent orientation versus only 22.9% of the non-looping pairs (Fig. 2C). We also observed that the motif match strength, as measured by the significance of a motif location to match the canonical CTCF motif [35], is higher for motifs involved in looping interactions (Fig. 2D). Together, the linear genome encodes several features, such as motif strength, orientation, and distance, that correlate with chromatin looping and can be used to predict such interactions.

### Chromatin loop prediction using 7C

To make use of both the condition specific ChIP-seq signals and the genomic features of CTCF motifs to predict chromatin loops, we trained a prediction model that takes only ChIP-seq data as input. To this end, we built a logistic regression model that takes into account only four features: the correlation coefficient between the ChIP-seq signals of the paired CTCF motifs (in a window of 1000 bp around the motif), the genomic distance between motifs, the orientation, and the (minimum) motif hit significance score (see Methods for details). For each ChIP-seq data set, we trained and evaluated a separate model (Fig. S2A). The method is implemented as the R package ‘sevenC’, which predicts chromatin loops using as only input a bigWig file from a single ChIP-seq experiment.

### Prediction performance evaluation

We used 10-fold cross-validation to assess the performance of the predictions on independent data that was not seen in the training phase. For each cutoff on the predicted interaction probability score, we computed the sensitivity, specificity and precision to plot receiver operator characteristic (ROC) and precision recall curves (PRC). Since only 4.2% of CTCF pairs are measured to interact, we mainly used the area under the PRC (auPRC) to evaluate prediction performance since, compared to ROC, the PRC gives a more accurate classification performance in imbalanced datasets in which the number of negatives outweighs the number of positives significantly [36]. Furthermore, we defined an optimal cutoff for the prediction probability *p* based on optimizing the f1-score. The six selected TF ChIP-seq data sets have optimal f1-scores at about *p* = 0.15 (Fig. S2B). For binary prediction, we provide a default prediction score threshold as the average of thresholds with optimal f1-score for the 10 best performing TF ChIP-seq datasets.

### Prediction performance of sequence features and 7C with single and multiple TF ChIP-seq data sets

First, we evaluated how the sequence-encoded features can predict chromatin interactions. For this, we built regression models that use only these features. Each of these features alone, CTCF motif hit significance, motif orientation or distance, were very poor predictors, and resulted in auROC between 0.67 and 0.74 (Fig. 3A) and auPRC scores between 0.08 and 0.09 (Fig. 3C). Using the three sequence features together improved prediction performance (auROC = 0.85, auPRC = 0.22).

**Fig. 3.**
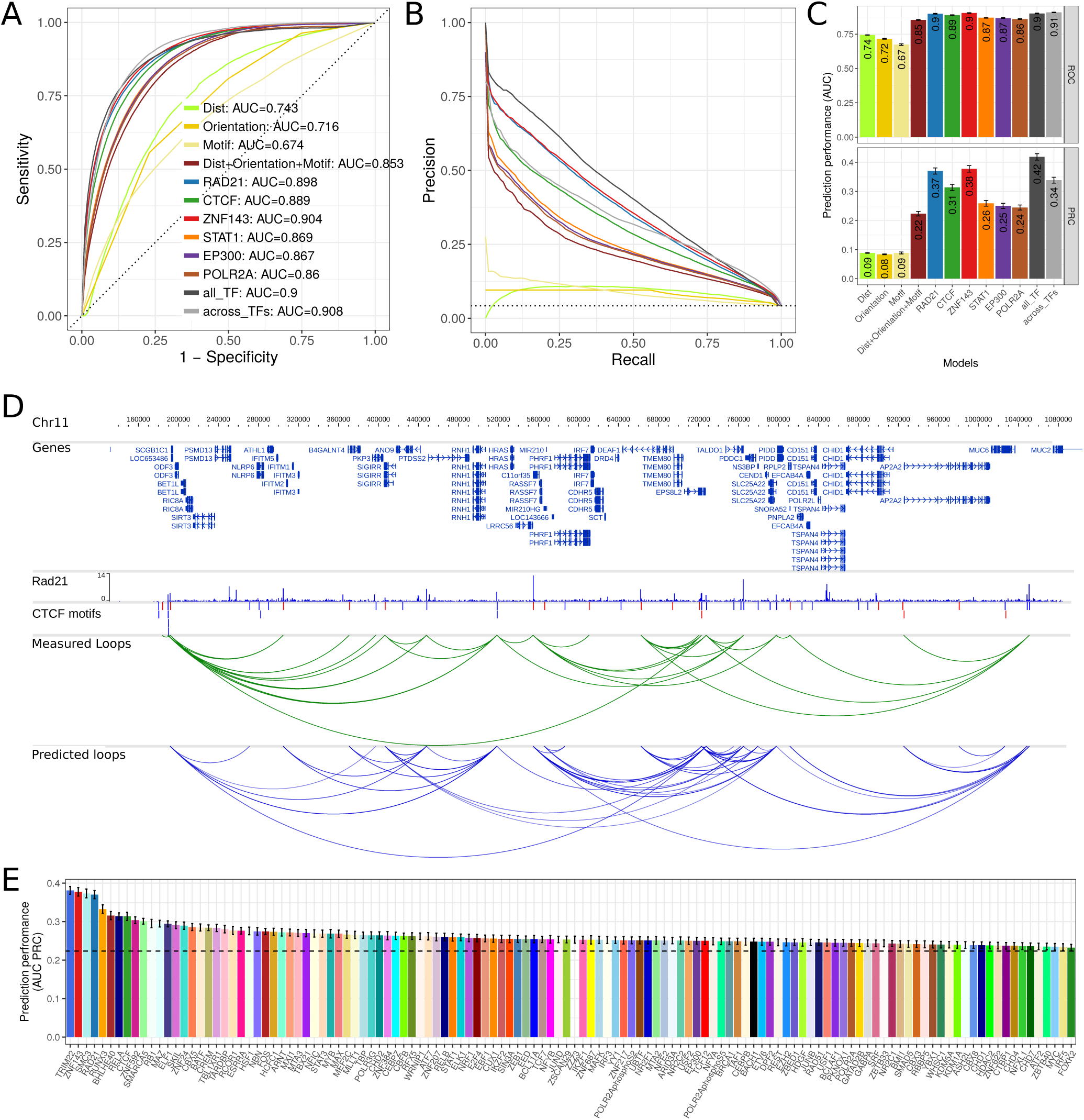
Prediction performance using cross-validation. **(A)** ROC plot for different models to predict chromatin looping interactions. The sensitivity (y-axis) is shown against the false discovery rate (1 – specificity, x-axis) for thresholds of the prediction score. Curves show averages of 10-fold cross-validation experiments. The models “Dist”, “Orientation”, and “Motif” contain only a single feature as indicated and all three genomic features are combined in the model “Dist+Orientation+Motif”. The models “RAD21”, CTCF”, “ZNF143”, “STAT1”, “EP300”, and “POLR2A”, contain the genomic features and the ChIP-seq correlation of the indicated factor. The model “all_TF” contains the genomic features and correlation of all indicated TFs. The model “across_TFs” contains the genomic features and a single correlation feature across the six ChIP-seq datasets as described in the main text. **(B)** PRC plot of precision against the recall for different prediction models. Color code as in (A). **(C)** Values of the area under the ROC (top) and PRC curves (bottom) as prediction performance. Error bars indicate standard deviation in 10-fold cross-validation experiments. **(D)** Example region on chromosome 11 in the genome browser showing: human genes, RAD21 ChIP-seq data in GM12878, CTCF motifs, CTCF motif pairs with that interact according to Hi-C or ChIA-PET data (green arcs) and predicted chromatin loops from RAD21 ChIPseq data using 7C (blue arcs). **(E)** Prediction performance of 7C as auPRC values for models with 124 TF ChIP-seq data sets from ENCODE. Error bars as in. The dotted horizontal line shows prediction performance of only the combined genomic features.

Next, we tested the addition of ChIP-seq data as feature in the prediction model using ChIP-seq data for each of six different TFs. Three of them, CTCF, RAD21, and ZNF143, have known function in chromatin loop formation [1,37–39], while STAT1, P300, and POL2, are to our knowledge not directly involved in chromatin loop formation (Fig. 1D). Adding any of these TF ChIP-seq datasets to the model increased prediction performance. STAT1, EP300, and POL2 only moderately increased prediction performance with auROC values between 0.86 and 0.87 (Fig. 3A) and auPRC between 0.24 and 0.26 (Fig. 3B, C). However, ChIP-seq of the known architectural proteins CTCF, RAD21, and ZNF143 resulted in markedly increased prediction performance with auPRCs of 0.31, 0.37, and 0.38 for CTCF, RAD21, and ZNF143, respectively (Fig. 3B, C). To test how much the performance depend on the actual truth set of measured loops, we trained and validated 7C on each individual Hi-C and ChIA-PET data set, as well as their intersection, and observed similar performance across data sets (Fig. S4A). For visual comparison, we show the actual looping interactions and 7C predictions on example region at chromosome 11 (Fig. 3D) and an overlay of 7C predicted loops with a Hi-C interaction heatmap (Fig. S3).

Next, we built a full model using the sequence based features and the ChIP-seq data of all six selected TFs. This only resulted in a slight increase of prediction performance to auPRC = 0.42 (Fig. 3B, C), indicating that a single ChIP-seq experiment might be sufficient for accurate prediction of chromatin loops. We also tested if a single value of correlation of ChIP-seq signal at both loop anchors across the six different TFs is predictive. Indeed, we find high prediction performance of auPRC = 0.34 for this approach. However, this was lower than using the correlation from single TF ChIP-seq experiments for RAD21 or ZNF143 and has the disadvantage of relying on ChIP-seq data from multiple experiments.

Another recently published method uses CTCF ChIP-seq peak heights together with motif orientation and distance in an iterative algorithm to predict chromatin interactions [40]. However, independent of the TF used, 7C yields higher specificity, precision and overall accuracy than this previously published method (Fig. S4B).

Together, these results show that sequenced based features alone have only a limited loop prediction performance, but integrating them with a single ChIP-seq experiment, 7C can predict chromatin loops with higher accuracy.

### Comparison of transcription factors by prediction performance

Our results can be used to better understand the molecular mechanisms of chromatin loop formation. We hypothesize that TFs whose ChIP-seq provides high prediction performance are likely to be functionally involved in chromatin looping. These TFs would be therefore interesting targets for further investigation of their potential function in chromatin looping.

To investigate this for as many TFs as possible, we used all available 124 TF ChIP-seq datasets from ENCODE for the human cell line GM12878 and compared transcription factors by their prediction performance. Notably, nearly all TF ChIP-seq data sets could increase the prediction performance of sequence-based features alone (Fig. 3E). However, there was a large variance in performance between TFs and a subset of TFs with high predictive power could be identified. These include for example the known architectural proteins mentioned above, CTCF, cohesin (RAD21 and SMC3), and ZNF143, but also factors, such as TRIM22, RUNX3, BHLHE40, or RELA, which might be interesting candidate factors with functional roles in chromatin loop formation.

### Prediction performance in other cell types and for different TFs

Next, we wanted to test if 7C is general enough to predict looping interactions in a cell type different to the one used to train it. To test this, we used the models presented above (trained with data from human GM12878 cells) to predict loops using as input ChIP-seq data from human HeLa cells. The prediction performance was assessed using as positives 12,480 loops (1.74 % of all motif pairs) identified in HeLa cells [13,16]. While prediction performance in HeLa cells is slightly lower as compared to the cross-validation in GM12878 cells, we see overall high prediction performance also in HeLa cells by ROC curves (auPRC up to 0.91, Fig. 4A) and PRC curves (auPRC up to 0.27, Fig. 4B,C).

**Fig. 4.**
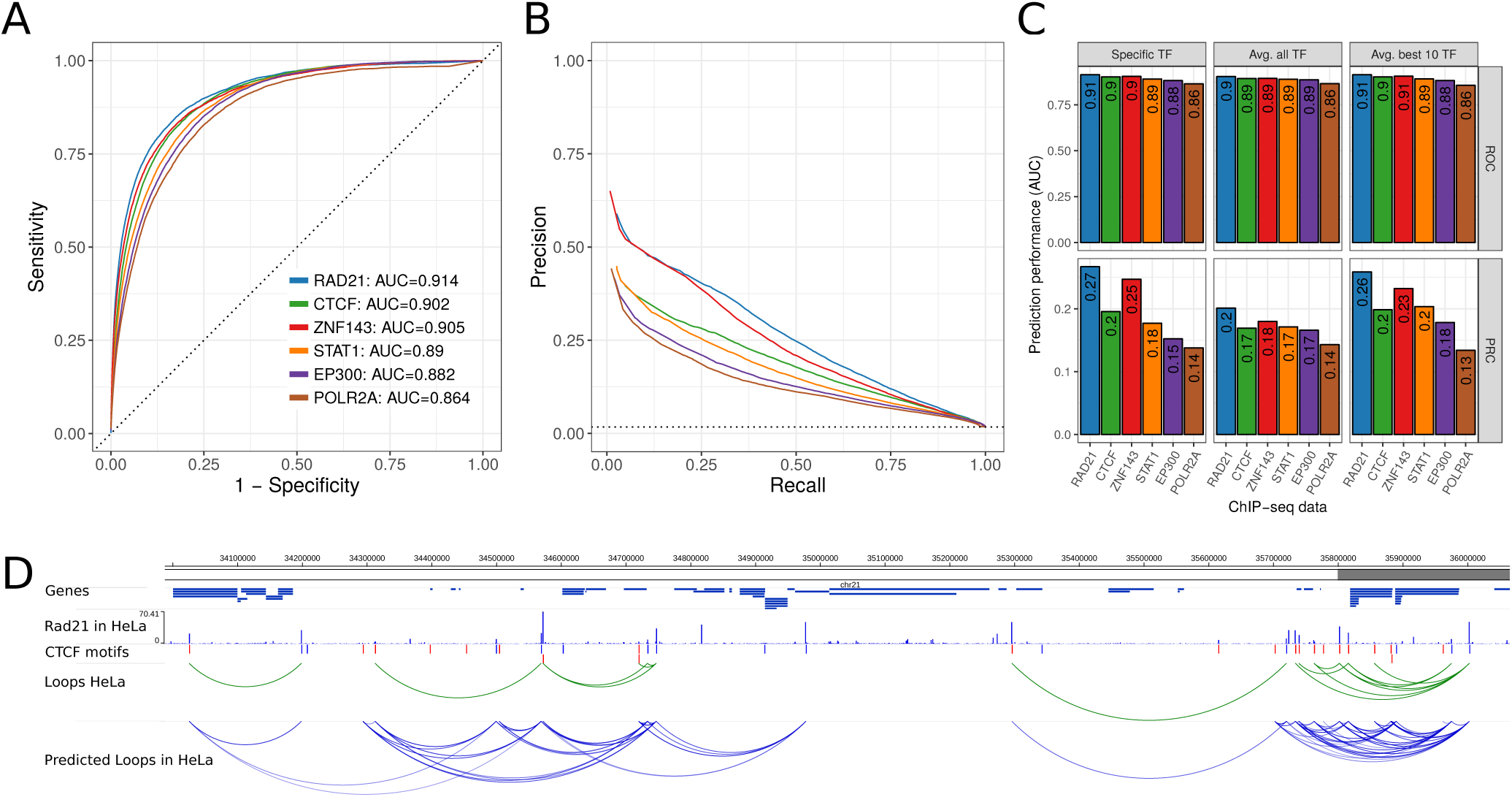
Prediction performance in HeLa cells using 7C trained in GM12878 cells. **(A)** ROC curve of prediction performance of six selected TF ChIP-seq data sets. The 7C model was trained using ChIP-seq and true loop data in human GM1287 but loops were predicted using ChIP-seq data of the same TFs in HeLa cells and true loop data in HeLa cells. **(B)** Precision-Recall curves for the same analysis as in (A). **(C)** Prediction performance as auPRC (top) and auROC (bottom) in HeLa for the six TF ChIP-seq data sets (x-axis) and 7C models trained for the specific TF (left), 7C with parameters averaged across all 124 TF models (center), and 7C with parameters as average of the 10 best performing TF ChIP-seq data sets (right). **(D)** Example region on human chromosome 21 with genes, RAD21 ChIP-seq data in HeLa, CTCF motifs, true loops in HeLa cells according to Hi-C and ChIA-PET (green arcs) and predicted chromatin loops from RAD21 ChIP-seq data in HeLa (blue arcs).

In this analysis, we compared the prediction performance of each specific TF model. However, in a real use case, one might not be able to train the model for a specific TF of interest and the model should predict loops for TFs that were not used in the training. Therefore, we built default 7C models by either averaging model parameters from all 124 TF models or by averaging across the model parameters of only the 10 best performing TFs. While all three approaches result in good prediction performance for the six selected TFs (Fig. 4C), the model averaging parameters across all TFs performs poorer than the ones of only the best 10 models, which are nearly as good as the specific TF models. This is consistent with similar results from cross-validation analysis in GM12878 data (Fig. S4C). Furthermore, we visually inspected chromatin loop predictions from RAD21 ChIP-seq data in HeLa at an example loci on chromosome 21 (Fig. 4D). In summary, these results show that 7C can predict chromatin looping interactions in different cell types that were not used to train it. Similarly, the 7C default prediction model performs nearly as good as a TF specific model. This makes 7C applicable for ChIP-seq data from diverse TFs in many different cell types and conditions.

### The high resolution of ChIP-nexus improves prediction performance

We wondered if other genomic measurements along the linear genome could provide similar signals at loop anchors potentially indicating looping interactions. Therefore, we used different genomic assays, such as DNase hypersensitivity (DNase-seq), ChIP-nexus and only ChIP-seq input control as input to our prediction methods (Fig. 5). Furthermore, we compared different signal types of ChIP-seq. During computational processing of ChIP-seq raw data, reads are shifted in 5’ direction by the estimated average fragment size [41,42]. The coverage of these shifted reads is then compared to coverage of input control experiment (fold change over control). Furthermore, a recent study quantified read pairs (qfrags) in a specific distance to each other as estimate for the actual fragment numbers detected by ChIP-seq [42]. For most of the TFs tested here, we observed that the ChIP-seq signal types ‘shifted reads’ and ‘qfrag’ have better loop prediction performance than the ‘fold change over control’ (Fig. 5). Interestingly, even the combination of ChIP-seq control with sequence features improves the prediction performance over using sequence features alone, indicating that cross-linking efficiency and density of chromatin itself is specifically distributed at chromatin loop anchors (Fig. 5). Also, DNase-seq, which measures chromatin accessibility, predicts looping interactions with similar accuracy than ChIP-seq input control (Fig. 5). This is consistent with specific open-chromatin profiles at TF binding sites [43,44].

**Fig. 5.**
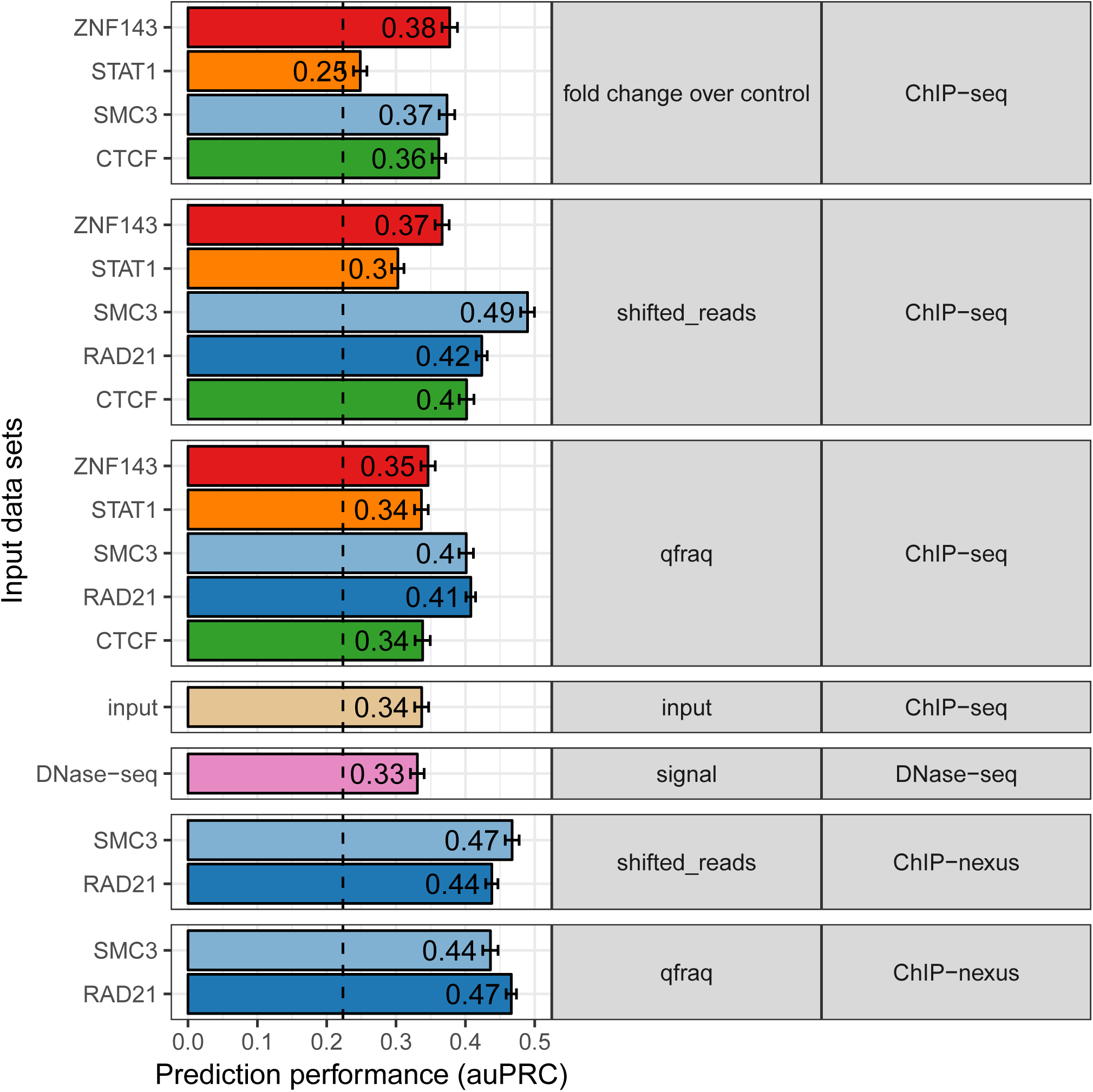
Higher resolution of ChIP-exo and ChIP-nexus improves prediction performance. Prediction performance as area under the precision recall curve (auPRC, x-axis) for 7C models with sequence features and different input data sets to predict chromatin looping (y-axis). Input data sets are grouped by signal-type (middle panel) and assay-type (right panel) and colored according to the TF (if any) used in the experiment.

However, using ChIP-nexus data for RAD21 and SMC3 [16], we could markedly improve chromatin loop predictions using 7C (Fig. 5). ChIP-nexus and ChIP-exo are variations of the ChIP-seq protocol, in which additionally, an exonuclease digestion step is applied to trim the DNA from the 5’ end until the actual bound protein [45,46]. These signals result in high-resolution binding footprints that can be used to identify different TF binding modes and cooperation with co-factors [34]. Therefore, we conclude, that the high-resolution binding profiles from ChIP-nexus allow to compute a more predictive binding signal similarity at chromatin loop anchors.

In summary, the comparison of different genomic signal types shows that cross-linking effect and chromatin density at chromatin anchors are predictive signals for long-range chromatin interactions and higher resolution TF binding assays, such as ChIP-nexus, result in improved prediction performance.

## Discussion

We have developed 7C to reuse ChIP-seq data, profiling the interactions of proteins with genomes, for the prediction of chromatin looping interactions between CTCF motif pairs within 1 Mb. We present this method as an alternative to dedicated techniques like Hi-C that directly measure genomic contacts. Since the results of ChIP-seq experiments are increasingly available for a large number of proteins, species, tissues, cell types, and conditions, our method offers an alternative when Hi-C data is not available, or cannot be produced due to cost or material limitations. Another major advantage of 7C over Hi-C is that the predictions are at a base pair resolution due to the use of paired CTCF motifs, while Hi-C only reaches resolutions of at best kilo base pairs at a high cost.

7C can use ChIP-seq data from any TF. It does not make any assumptions about the mode of binding or function of the TF and the TF does not need to be involved in the loop formation. It is enough if the binding of the TF close to a CTCF-based loop results in symmetrical ChIP-seq traces. Regardless, it is possible that TFs binding very close and very often to the anchors of loops will be better predictors, and TFs with functions in loops might as well have stronger and more reproducible binding there. For example, it is interesting to note that ZNF143 has been known for a while to bind close to CTCF-cohesin in the anchors of chromatin loops (e.g. [7]), and very recently it has been reported as a regulator of loop formation [47]. Using the HIPPIE database [48], we found that while there is no experimental evidence of direct interaction between CTCF and ZNF143, there is one single protein that is reported to interact with both CTCF and ZNF143, the chromodomain helicase DNA binding protein 8 (CHD8; see [49] and [50], respectively). So while no direct interaction between ZNF143 and CTCH is known, they could form a complex via CHD8. Differently, for another very good predicting factor such as TRIM22 no connection to loop architecture has been yet established; they might be none, or it could occur in an unexplored condition. TRIM22 is an antiviral protein whose expression is triggered by interferon, with cytoplasmic and nuclear localizations, and a variety of functions (see e.g. [51]); we would be careful to suggest that this protein might have yet another function controlling genomic structure.

Other computational approaches were developed to predict genomic contacts or assign regulatory regions to target genes. A commonly used approach is to compare activity signals at enhancers and promoters across many different conditions or tissues [52–56]: high correlation indicates association and potential physical interactions between enhancers and genes. However, these approaches lose the tissue specificity of the interactions. Other approaches integrate many diverse chromatin signals such as post-translational histone modifications, chromatin accessibility, or transcriptional activity [57–62], and combine them with sequence features [63], or evolutionary constrains [64]. While these methods predict enhancer-gene association with good performance, they require for each specific condition of interest a multiplicity of input datasets, which are often not available.

Further computational approaches try to directly predict chromatin interactions by using diverse sequence features [65] or multiple chromatin features such as histone modifications [66–68] or transcription [69]. One study makes use of the more recently discovered CTCF motif directionality to predict loop interactions from CTCF ChIP-seq peak locations [40], but has lower prediction performance than 7C (Fig. S4B) Another study combines CTCF binding locations and motif orientation with polymer modeling to predict Hi-C interaction maps [29]. However, none of these studies predicts chromatin loops from ChIP-seq signals of TFs different from CTCF by taking the CTCF motif orientation into account. Furthermore, CTCF binding sites are often only considered, when the signal is strong enough for peak calling algorithms to identify binding sites. In contrast, 7C takes the distribution of ChIP-seq signals from all TFs into account without a peak-calling step. Furthermore, the other studies, except one [40], do not provide a tool for the direct prediction of pairwise interactions from single ChIP-seq experiments. Interestingly, shadow peaks in ChIP-seq data of insulator proteins in *Drosophila* were previously associated to long-range interactions [70] and used to study the contribution of sequence motifs and co-factors in loop formation [71], but not to directly predict chromatin loop interactions.

Compared to other predictive methods mentioned above, our approach has the advantage to directly predict chromatin looping interactions, and not enhancer-promoter associations, by making use of ChIP-seq signals from a single experiment with respect to CTCF motifs. However, many enhancer-promoter interactions occur in the span of interacting CTCF binding sites, which were described to form insulated neighborhoods [72,73]. Therefore, 7C predictions can help to associate enhancers to genes when they are located between predicted loop anchors. The motifs give the prediction a base pair resolution. In fact, given several CTCF motifs within a 1kb genomic bin, our looping prediction approach can be used to decide which of the CTCF sites is actually involved in the measured interactions and thus increase resolution even when Hi-C data is available. We showed that our approach, 7C, can work with just a single ChIP-seq experiment for many different TFs, making it usable for many diverse conditions of interest. Therefore, 7C can be used complementary to existing enhancer-promoter association tools or can be integrated in such predictive models to improve them.

Currently, our method, by using CTCF motifs, focuses on CTCF mediated chromatin loops. It is very likely that other DNA binding proteins mediate loops: for example, recent studies suggest that other TFs are involved in enhancer promoter interactions during differentiation [74] and knockout of transcriptional repressor YY1 and other candidate factors result in loss of chromatin loops [75]. Using motifs predicted for these different transcription factors, or combinations thereof, are open avenues for the future extension of our method.

## Conclusion

We demonstrated that TF binding signals of ChIP-seq experiments at CTCF motifs are predictive for chromatin looping. We provided a method, 7C, that is simple to use and integrates these signals with genomic sequence features to predict long-range chromatin contacts from single ChIP-seq experiments. 7C is freely available as R/Bioconductor package (bioconductor.org/packages/sevenC). The analysis of ChIP-seq experiments for 124 different TFs highlighted the role of cohesin, ZNF143 and CTCF in chromatin loop formation, but also suggested many other TFs, such as TRIM22, RUNX3, and BHLHE40, to be functionally involved in chromatin looping, likely in cooperation and protein interaction (direct or indirect) with CTCF at loop anchors.

Since our method needs only a single ChIP-seq experiment as input, it enables the analysis of chromatin interactions in diverse cell types and conditions, where Hi-C like data is not available. Therefore, 7C can be used to enable condition specific associations of distal TF binding sites and enhancers to promoters of target genes. These might allow the interpretation of non-coding genetic variants by genes in physical contact with the variant loci in a specific cell type or condition of interest. Furthermore, 7C might improve the resolution of Hi-C interaction maps by facilitating base-pair specific pairing of CTCF motifs located in bins of several kb. With these applications, 7C increases the value of ChIP-seq datasets, which now can be used to improve the analysis of 3D genome folding and their dynamic changes between diverse cell types and conditions.

## Methods

### CTCF motifs in the human genome

The recognition motif of CTCF is well defined and available from the JASPAR database (MA0139.1) [76]. We downloaded TF binding site predictions with the CTCF motif (MA0139.1) in the human genome hg19 from the JASPAR database (http://expdata.cmmt.ubc.ca/JASPAR/downloads/UCSC_tracks/2018/hg19/tsv/MA0139.1.tsv.gz). Motif hits were filtered for p-value ≤ 2.5 x 10^-6^, resulting in 38,316 highly significant CTCF motif hits genome-wide and 717,137 motif pairs within 1 Mb genomic distance that are considered as potential loop interaction anchors in this study.

### Loop interaction data for training and validation

For training and validating the prediction model we used 9,448 published loops derived from high-resolution *in-situ* Hi-C experiments [13] and 206,399 CTCF and Pol2 ChIA-PET interactions [16] in human GM12878 cells. We considered each CTCF motif pair as positive (true looping interaction) if there was at least one measured looping interaction for which each loop anchor overlapped one of the CTCF motifs. Overlaps were calculated using the R package *InteractionSet* [77]. This resulted in 30,025 (4.2%) of 717,137 candidate motif pairs that were labeled as true looping interactions in GM12878. For the prediction validation in HeLa cells we used the 3,094 Hi-C loops and 402,722 ChIA-PET interactions for CTCF and Pol2 in HeLa from the same studies [13,16] and labeled 12,480 (1.7 %) of motif pairs as true loops in HeLa cells.

### ChIP-seq datasets in GM12878 cells

We downloaded publicly available ChIP-seq data from the ENCODE data portal [21,22] by requiring the assay to be ChIP-seq, the target to be a transcription factor, the biosample term name to be GM12878, the genome assembly to be hg19, and the file-type to be bigWig. Furthermore, we filtered the data to have output type ‘fold change over control’ or ‘signal’ and to be built from two replicates. Then we selected for each TF only one unique experiment as bigWig file with either output type ‘fold change over control’ or, if unavailable, output type ‘signal’. This resulted in 124 ChIP-seq experiments for different TFs (Table S1). ChIP-seq data for HeLa were retrieved analogously and filtered for the selected targets: RAD21, CTCF, ZNF143, STAT1, EP300, and ZNF143 (Table S2).

### ChIP-seq data types

To analyze the effect of different ChIP-seq signal types and other genomic assays on loop prediction performance, we selected five TFs (ZNF143, STAT1, SMC3, RAD21, and CTCF) and downloaded the mapped reads of ChIP-seq experiments as BAM files from the ENCODE data portal [22] and from the UCSC Genome Browser [78]. Furthermore, we downloaded signal tracks as bigWig files for ChIP-seq input control experiment and DNase-seq experiments in GM12878 cells. File accession identifiers and download links are provided in Table S3. We used the ChIP-seq peak caller *Q* [42] with option ‘-w’ for each human chromosome to generate signal tracks in BED format of shifted reads and qfrags. ‘Shifted reads’ are counts of mapped reads that are shifted in 5’ direction by half of the estimated fragment size. ‘qfrags’ are pairs of forward and reverse mapped reads within a given distance [42] and are shown to improve signal to noise ratio in ChIP-seq peak calling [42]. We then combined resulting BED files from all chromosomes and converted them to the bedGraph and bigWig formats using the *bedtools* [79] and *bedGrpahtoBigWig* tools from the UCSC Genome Browser [80].

### ChIP-nexus data processing for RAD21 and SCM3

ChIP-nexus data for RAD21 and SMC3 in GM12878 cells were published recently [16]. We downloaded the corresponding raw reads from the Sequence Read Archive (SRA) (Run IDs SRR2312570 and SRR2312571). Reads were processed using *felxcut* for barcode removal and adapter trimming as recommended in the user guide of the *Q-nexus* tool [81]. Reads were than mapped to human genome hg19 using *Bowtie* version 2.3.2 with default settings. Duplicate reads were removed using *nexcat* [81]. Finally, we created shifted-reads and qfraq profiles using *Q-nexus* [81] with options ‘--nexus-mode’ and ‘-w’ for each chromosome and combined them to bigWig files as described above.

### Similarity of ChIP-seq profiles as correlation of coverage around motifs

For each CTCF sequence motif in the human genome, we quantified the number of reads overlapping each base within +/- 500 bp around the motif center. This results in a vector *x*_*i*_=(*x*_*i*, 1_, *x*_*i*, 2_,…, *x*_*i, n*_) where *x*_*i,k*_ is the ChIP-seq signal at position *k* around CTCF motif *i*. ChIP-seq signal vectors for motif hits reported on the minus strand were reversed because CTCF motif sites are assumed to be symmetrically aligned to each other when cooperating at loop anchors (Fig. 1A) [13,16,27–29]. For all considered pairs of CTCF motifs *i* and *j*, we calculated the ChIP-seq profile similarity as Pearson correlation coefficient *ri, j* of the corresponding coverage vectors *x*_*i*_ and *x* _*j*_.

### Genomic sequence features of chromatin loops

Besides the correlation of ChIP-seq profiles, we used genomic features of motif pairs as features to predict interactions. The distance *d* is the number of bp between the two motif centers. The categorical variable orientation *o* is either, *convergent, forward, reverse*, or *divergent*, depending on the orientation of CTCF motifs in the pair (+-, ++, –, and −+, respectively). The motif hit similarity *s* is the minimum of the two motif hit scores in each pair; we derived these motif scores from the JASPAR motif hit tracks as −log10 transformed p-values [35].

### 7C prediction model

We used a logistic regression model to predict the log-likelihood probability of CTCF motif pairs to perform chromatin looping interactions. The probability *p* that two sites interact is modeled as:

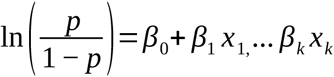

where *β* are the unknown model parameters and *x*_1,_ *… x*_*k*_ the features.

More specifically, for the 7C model with a single ChIP-seq experiment as input, the logistic regression model for the interaction probability *p* is:

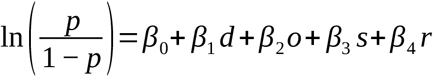

Parameters were estimated using the function ‘glm()’ with option ‘family=binomial()’ in R during model training as described below.

### Training and validation of prediction model

We used the R package *rsample* for 10-fold cross-validation. Thereby, we randomly split the dataset of CTCF motif pairs into ten equal sized subsets. For each round of cross-validation one subset is held out (test dataset) and the model parameters are trained on the remaining 90% of the samples (training dataset). The model parameters are shown for six selected TFs and combined models in Supplementary Figure S2A. For each split, the performance of the model is than evaluated on the test dataset. For prediction performance in HeLa cells, we trained on all motif pairs using ChIP-seq and true loops from GM12878 cells and evaluated performance on all motif pairs using the true loop data in HeLa.

### Analysis of prediction performance

We quantified prediction performance using the receiver operating characteristic (ROC) and precision recall curves (PRC) as implemented in the R package *precrec* [82].

Given the number of true positives (TP), true negatives (TN), false positives (FP), and false negatives (FN), the sensitivity is defined as TP/(TP+FN), specificity as TN/(TN+FP), precision as TP/(TP+FP), and recall as TP/(TP+FN). For each cross-validation split, the area under the curve is computed separately, and the mean across splits together with the standard deviation reported. To get binary prediction outputs, we computed the f1-score as harmonic mean of precision and recall for all prediction scores on all cross-validation folds using the R package *ROCR* [83]. Then we computed the prediction score that maximizes the f1-score as default cutoff for binary prediction output (Fig. S2B).

### Comparison to a previously published tool

We downloaded the script provided by Oti et al 2016 [40] from https://doi.org/10.5281/zenodo.29423. We downloaded peaks from the same CTCF ChIP-seq experiment that was analyzed with 7C from ENCODE (https://www.encodeproject.org/files/ENCFF710VEH/@@download/ENCFF710VEH.bed.gz). We filtered the CTCF motifs described above to overlap a peak region and assigned to each motif a score between 0 and 1000 according to the overlapping peak height. This data was then used as input to the script “ctcf_peaks2loops.py” to predict loops. We computed sensitivity, specificity, precision, and accuracy by overlap with the true loops described above and compared the performance to loop predictions of 7C.

### Implementation of 7C and compatibility to other tools

We implemented 7C as R package, termed *sevenC*, by using existing infrastructure for chromatin interaction data from the *interactionSet* package [77] and functionality for reading bigWig files from the *rtracklayer* package [84] from the Bioconductor project [85]. Predicted loops can be written as interaction tracks for visualization in the WashU Epigenome Browser [86] or as BEDPE format using the *GenomicInteractions* package [87] for visualization in the Juicebox tool [88]. The package is freely available and easy to install from Bioconductor https://bioconductor.org/packages/sevenC. All analysis presented in this work were implemented in R and all scripts used have been made available in a separate GitHub repository: https://github.com/ibn-salem/sevenC_analysis.

## Declarations

### Ethics approval and consent to participate

Not applicable

### Consent for publication

Not applicable

### Availability of data and material

The method 7C is implemented as R packages sevenC and can be downloaded with documentations from Bioconductor: http://bioconductor.org/packages/sevenC (DOI: <u>10.18129/B9.bioc.sevenC</u>). The source code is also available on GitHub: https://github.com/ibn-salem/sevenC. The source code for all analyses in this manuscript is available in a separate GitHub repository: https://github.com/ibn-salem/sevenC_analysis. All the genomic data analyzed here are freely available to be downloaded from the GEO repository or ENCODE as described in the methods section.

### Authors’ contributions

JI developed and implemented the method, conceived the study and performed all analyses. MA supervised the study. JI and MA wrote the manuscript.

## Acknowledgments

The authors thank all members of the CBDM group for fruitful discussions, especially Katerina Taškova for ideas on modeling and cross-validation. We further thank Idan Gabdank (Stanford University) for support in understanding ENCODE data set and updating it, Takaya Saito (University of Bergen) for updating and improving the *precrec* R package, Peter Hansen (Charité, Berlin) for support in ChIP-seq and ChIP-nexus analysis using *Q*, Morgane Thomas-Chollier (Ecole normale supérieure) for discussions on genome-wide motif analysis, and Sebastiaan Meijsing (Max Planck Institute for Molecular Genetics) for initial discussions on ChIP-exo profiles at chromatin loop anchors.

## Supplementary Tables

**Table S1**

Metadata of ChIP-seq experiments from ENCODE in human GM12878 cells with accession ID and download link.

**Table S2**

Metadata of ChIP-seq experiments from ENCODE human HeLa cells with accession ID and download link.

**Table S3**

Accession numbers and download URLs for data sets used in data type comparisons.

### Supplementary Figures

**Fig. S1**

**Hi-C and ChIA-PET interactions and their overlap with CTCF motif pairs (A)** Number of genome-wide CTCF motifs by motif hit significance cutoff. **(B)** Number of CTCF motif pairs within 1 Mb distance by motif hit significance. **(C)** Percent of CTCF motif pairs that overlap with experimentally measured Hi-C and ChIA-PET loops by the motif hit significance. **(D)** Upset plot of true loop data sets (rows) and their size (horizontal bars) with their intersections (columns, and vertical bars) based on the number of overlapping CTCF motif pairs. **(E)** Distribution of interaction span (distance between anchors) of Hi-C loops and ChIA-PET loops in GM12878 that are used as gold standard. The dotted red line indicate the distance cutoff (1Mb) used in this study. **(F)** Number and percent of Hi-C and ChIA-PET loops that overlap with CTCF motif pairs within a distance of 1 Mb. **(G)** Number and percent of Hi-C and ChIA-PET loops that overlap with 1, 2, 3, 4, 5 or more than 5 CTCF motif pairs. The percent values are relative to all loops that overlap at least one CTCF motif pair.

**Fig. S2**

**7C model parameters and optimal cut-offs for binary prediction. (A)** Parameter values of the logistic regression model in 7C for different features (columns), separated for different models (rows). Average of model parameters of model training in 10-fold cross-validation is shown with error bars indicating the standard deviations. While the first six rows represent the models with the indicated TF ChIP-seq data and the genomic features, “Avg. all TF” is the average across all 124 TFs analyzed and “Avg. best 10 TF” is the average across the best ten performing TF models. **(B)** Prediction performance as f1 score (y-axis) for different cutoffs on the prediction probability *p* for the six selected models.

**Fig. S3**

High resolution Hi-C map with 7C loop predictions. The red color intensity shows Hi-C interactions frequencies at an example locus of chromosome 1. The blue squares indicate 7C loop predictions using a Rad21 ChIP-seq experiment. The figure was created using the Juicebox tool by loading the combined Hi-C data set in GM12878 from [13] with mapping quality MAPQ ≥ 30 at a resolution of 5kb.

**Fig. S4**

**(A)** Prediction performance (auPRC) of 7C when trained and evaluated on different datasets of experimentally measured loops as gold standard. Rao_GM12878 refers to Hi-C loops from [13], Tang2015_GM12878_CTCF, and Tang2015_GM12878_RNAPII to ChIA-PET loops using CTCF or Polymerase II as the target [16]. In Union, all datasets were taken together, and in Intersection, only those CTCF motif pairs that were measured in all datasets were considered positive. **(B)** Prediction performance of 7C with six different TFs compared to the method by Oti et al. [40]. The figure shows from top to bottom the accuracy, precision, sensitivity, and specificity of the predictions. The Top **(C)** Prediction performance as auPRC (top) and auROC (bottom) of four different models (colors) on ChIP-seq data for six selected TFs (x-axis). ‘Specific TF’ is the model fitted using the ChIP-seq data indicated on the x-axis, ‘RAD21’ is the model trained on RAD21 ChIP-seq data, ‘Avg. all TF’ is a model averaged across all 124 models of analyzed TFs, and ‘Avg. best 10 TF’ is the averaged model across the 10 best performing models.

