## Supplementary figures and images for "Computational Chromosome Conformation Capture by Correlation of ChIP-seq at CTCF motifs"

### Supplemental Figure S1

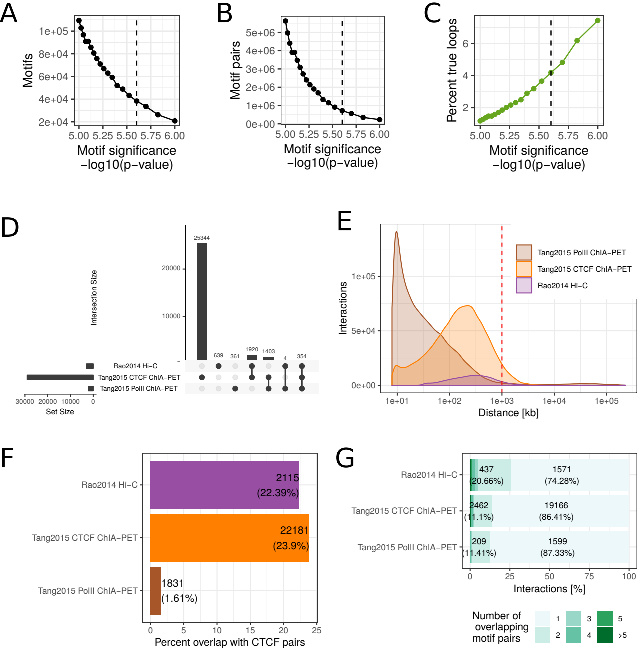

### Supplemental Figure S2

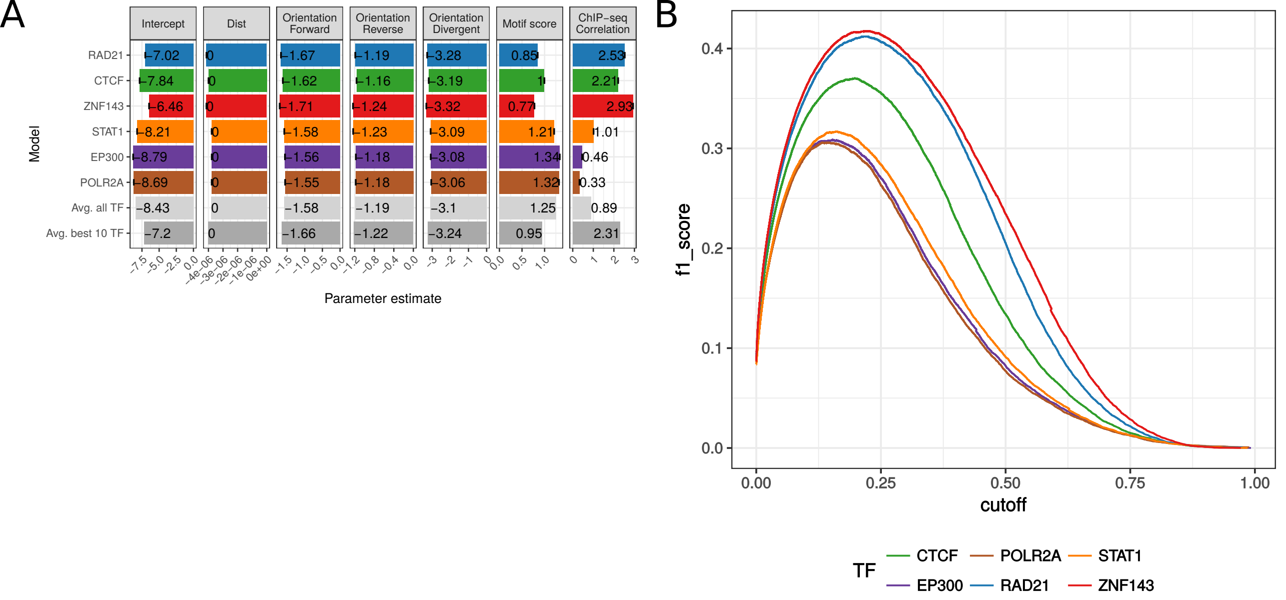

### Supplemental Figure S3

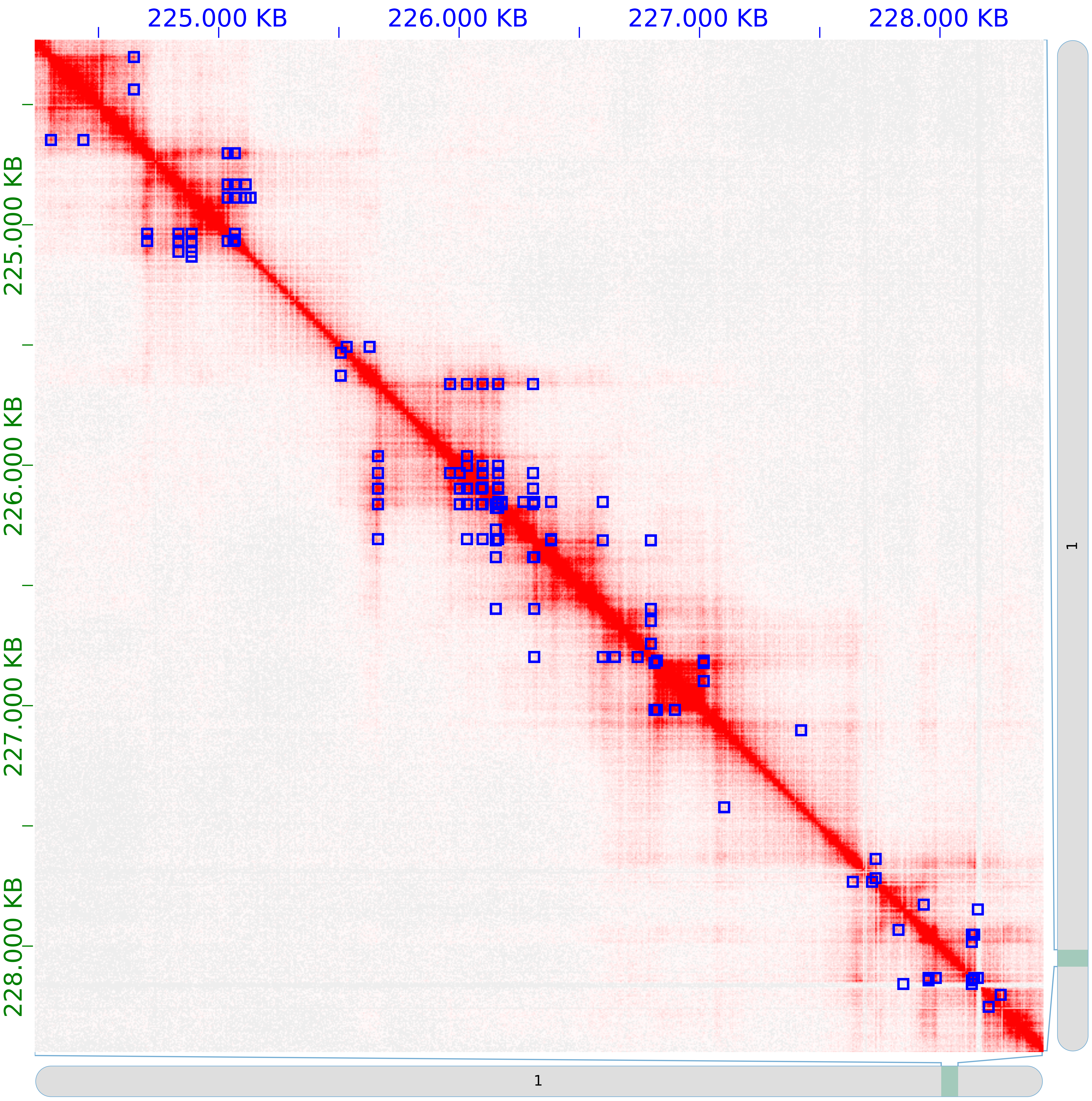

### Supplemental Figure S4

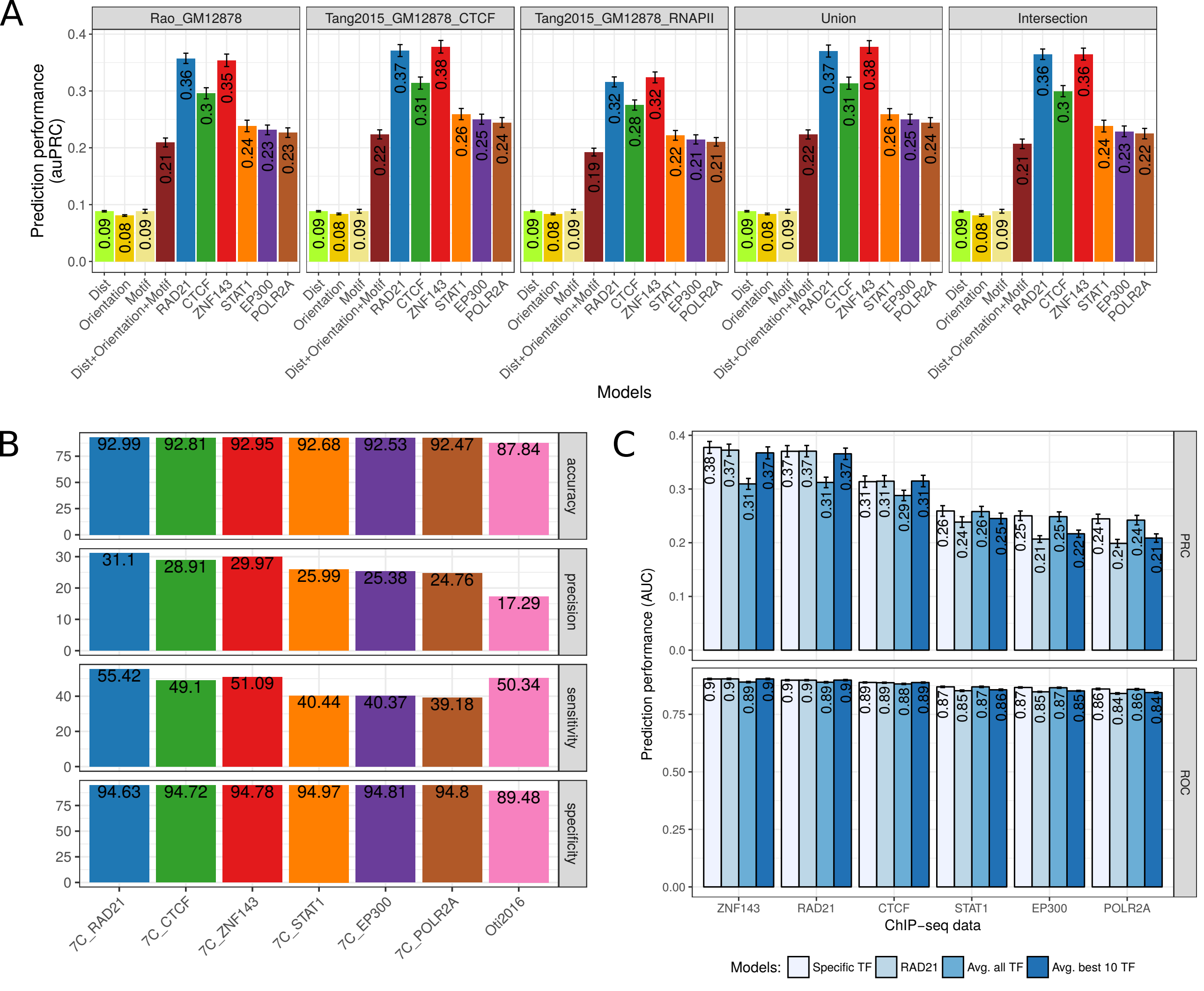
